# Transcriptome analyses of β-thalassemia -28 (A>G) mutation using isogenic cell models generated by CRISPR/Cas9 and asymmetric single-stranded oligodeoxynucleotides (assODN)

**DOI:** 10.1101/2020.06.18.159004

**Authors:** Jing Li, Ziheng Zhou, Hai-Xi Sun, Wenjie Ouyang, Guoyi Dong, Tianbin Liu, Lei Ge, Xiuqing Zhang, Chao Liu, Ying Gu

## Abstract

β-thalassemia, caused by mutations in the human hemoglobin (*HBB*) gene, is one of the most common genetic diseases in the world. *HBB* –28 (A>G) mutation is one of the five most common mutations in China patients with β-thalassemia. However, few studies have been conducted to understand how this mutation affects the expression of pathogenesis related genes including globin genes due to limited homologous clinical materials. Therefore, we first developed an efficient technique using CRISPR/Cas9 combined with asymmetric single-stranded oligodeoxynucleotides (assODN) to generate a K562 cell model of *HBB* −28 (A>G) named K562^−28 (A>G)^. Then, we systematically analyzed the differences between K562^−28 (A>G)^ and K562 at the transcriptome level by high-throughput RNA-seq pre- and post-erythrogenic differentiation. We found that *HBB* −28 (A>G) mutation not only disturbed the transcription of *HBB* but also decreased the expression of *HBG*, which may further aggravate the thalassemia phenotype and partially explain the severer clinical outcome of β-thalassemia patients with *HBB* −28 (A>G) mutation. Moreover, we found K562^−28 (A>G)^ cell line is more sensitive to hypoxia and showed a defective erythrogenic program compared with K562 before differentiation. In agreement, p38MAPK and ERK pathway are hyperactivated in K562^−28 (A>G)^ after differentiation. Importantly, all above mentioned abnormalities in K562^−28 (A>G)^ were reversed after correction of this mutation with CRISPR/Cas and assODN, confirming the specificity of these phenotypes. Overall, this is the first time to analyze the effects of the *HBB* - 28 (A>G) mutation at whole-transcriptome level based on isogenic cell lines, providing a landscape for further investigation of the mechanism of β-thalassemia with *HBB* −28 (A>G) mutation.

## Background

Beta-thalassemia is one of most common autosomal recessive blood diseases spread worldwide, caused by either point mutations or deletions in the β-globin (*HBB*) gene^[1, 2]^. Approximately 80 to 90 million people carried β thalassemia, and about 300,000 children with severe thalassemia are born every year^[3–5]^.Mutations or deletions of β-globin genes result in the reduction of hemoglobin β chain (β-globin), deformation of hemoglobin tetramer and subsequent lysis of erythrocytes, finally causing oxygen shortage, bone deformity, organ dysfunction and even organ failure in many parts of the human body^[6, 7]^. As for the thalassemia major patients, life-long blood transfusion and iron chelation treatments are required for survival, but often accompanied by numerous complications, including arrhythmia, congestive heart failure, hypothyroidism, hypoparathyroidism, hypogonadism, diabetes, osteoporosis, liver cirrhosis and recurrent infections. Thus, thalassemia has threatened millions of people’s lives for decades and is still a major public health issue^[8, 9]^.

β-thalassemia mutations are prevalent in Southern part of China and Southeast Asia, and *HBB* −28 (A>G) mutation is one of the five most common HBB mutations carried by β-thalassemia patients in China^[7, 10]^. In *HBB* −28 (A>G) mutation, adenine (A) base located at 28 base pairs upstream from the cap site is mutated to guanine (G), disrupting the binding of transcription factor of ATA box and decreasing the RNA expression of *HBB*^[11]^. Patients with homozygous or compound heterozygous −28 (A>G) mutation may develop severe anemia or intermedia anemia^[6, 11]^. Although the description of severe thalassemia has been first published over 90 years ago and a considerable amount work has been reported to refine the understanding of the pathophysiology of thalassemia syndromes in the past 50 years, the cellular and molecular basis of this group of diseases are still not thoroughly investigated ^[12]^. In particular, few studies have been conducted on *HBB* −28 (A>G) mutation to understand how this mutation affects gene expression at transcriptome level, although correcting the *HBB* −28 (A>G) mutation with base editing (BE) system has also been reported in human iPS cells and reconstituted embryos ^[7]^.Without full understanding of the defects at molecular level, especially in the short and long run, it will be difficult to comprehensively evaluate the rescue effect after changing the mutation back. In a recent study, high-throughput RNA-sequencing has been used to compare control samples with patient samples carrying a novel *HBB* mutation (*HBB*: c.51C>T). It shows that hemopoiesis, heme biosynthesis, response to oxidative stress and other cellular activities pathway were directly or indirectly enriched by differentially expressed genes related to β-thalassemia ^[13]^, suggesting genome-wide RNA-seq analysis is a useful approach to understand the mechanism of β-thalassemia with different mutations. However, control samples in this study are allogenic, and different genetic backgrounds and mixture of short and longterm effect would prevent deep understanding of effect of each mutation.

In order to explore the specific impact of *HBB* −28 (A>G) mutation on erythroid differentiation and how it affects genome-wide gene expression without confounding factors, such as comparison of allogenic samples, we used CRISPR/Cas9 gene-editing system combined with asymmetric single-stranded oligodeoxynucleotides (assODN) to generate the disease model of isogenic K562 cell lines ^[14, 15]^, and then conducted transcriptome analysis by high-throughput RNA-sequencing. The mutant cell line was derived from immortalized K562 and named as K562^−28 (A>G)^, showing no detectable *HBB* gene expression and enabling comparative studies in the same genome background. We found that *HBB* was prevented from transcriptional expression, and GO and KEGG analysis revealed that K562^−28 (A>G)^ cell line is more sensitivity to hypoxia and present a defective erythrogenic program when compared with wild type K562 before erythroid differentiation. In agreement, p38MAPK and ERK pathway are hyperactivated in K562^−28 (A>G)^ after differentiation. Interestingly, *HGB* showed a lower rate of induction in K562^−28 (A>G)^ when compared with wild type K562 after erythroid differentiation. Taken together, our study is the first time to analyze the effects of the *HBB* −28 (A>G) mutation at whole-transcriptome level based on isogenic cell lines. The unraveled molecular biomarkers and signaling pathways that affected in K562^−28 (A>G)^ cell line may be further investigated as therapeutic targets to improve the quality of life for those β-thalassemia patients in future studies.

## Results

### Generation of *HBB-28* (A>G) mutant cell line by CRISPR/Cas9 and asymmetric single-stranded oligodeoxynucleotides

To study how *HBB* −28 (A>G) mutation affects gene expression at the genome-wide level, isogenic human cell line carrying this mutation was generated. The diagram of generating mutant cell line was shown, including transfection, CRISPR/Cas editing, single cell sorting, cell characterization and expansion (Fig. 1A). As previous studies demonstrated that using single-strand oligodeoxynucleotides (ssODNs) ^[16]^ or asymmetric double-strand DNA ^[17]^ as repair template resulted in higher efficiency of accurate replacement of target sequences through homology directed repair (HDR), we developed a new technology to combine CRISPR/Cas with asymmetric ssODNs (assODNs). To generate *HBB* −28 (G>A) mutation, sgRNA mediating DNA cleavage 3bp aside from the −28 mutation site and assODN with 36bp on the PAM-distal side and 91bp on PAM-proximal side of the cutting site were used (Fig. 1B). K562, a human erythroleukemia line that resembles undifferentiated erythrocytes, was transduced with these Cas/sgRNA and assODNs. As expected, we identified one cell line with homozygous mutation of *HBB* −28 (A>G) by Sanger sequencing and named it as K562^−28 (A>G)^ (Fig. 1C). In order to reconfirm the functional mutation of *HBB* −28(A>G), we used qPCR and ELISA to detect the expression of *HBB* mRNA and HBB protein respectively. In agreement with the sequencing result, the expression of *HBB* mRNA and HBB protein was undetectable in K562^−28 (A>G)^ cell line. In contrast, wild-type K562 showed a considerable expression of *HBB* mRNA and HBB protein even without erythroid differentiation (Fig. 1D and 1E). These results suggested that the mutant cell line of *HBB* −28(A>G) was successfully generated, in which the expression of *HBB* was eliminated.

**Figure 1.**
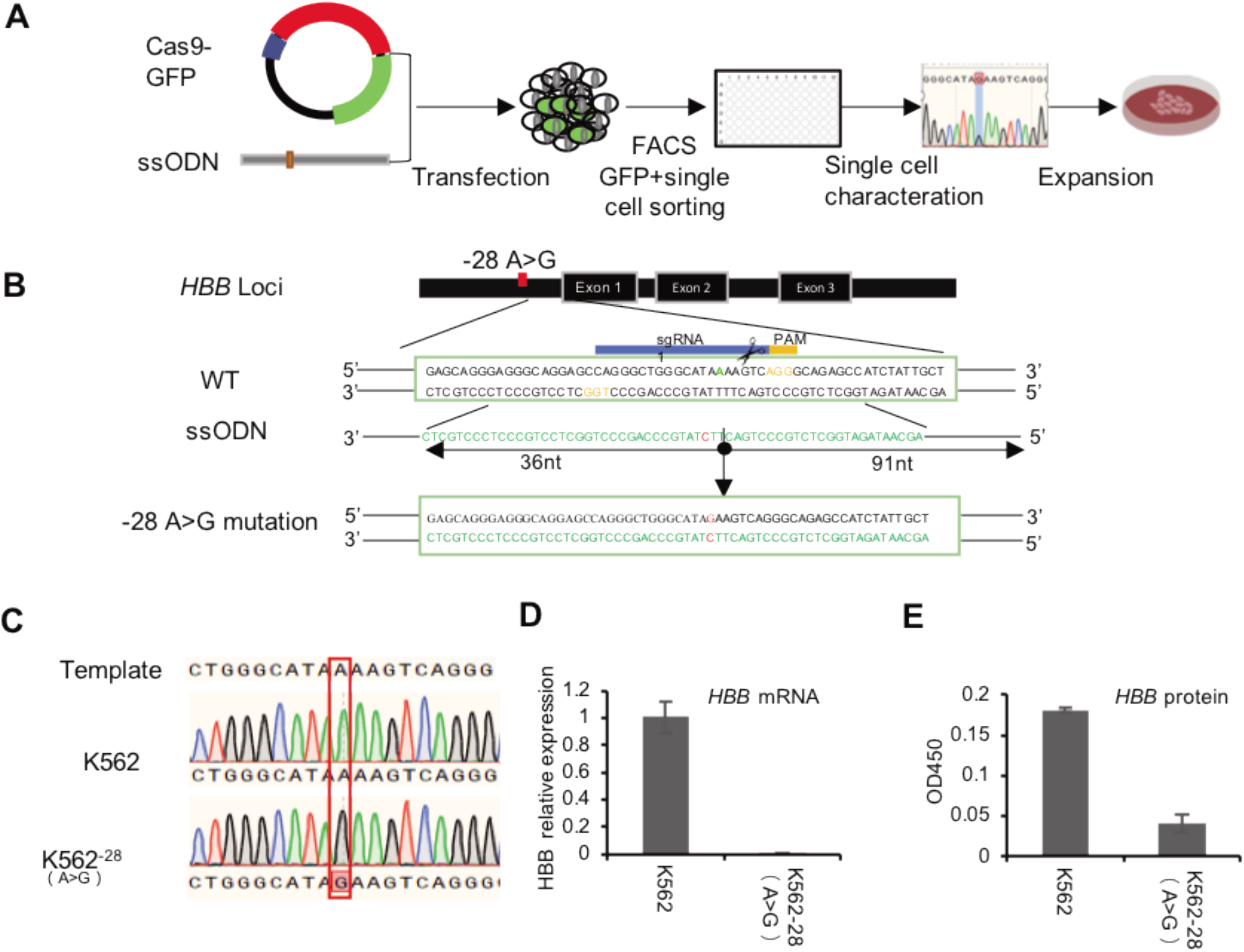
Generation of K562^−28 (A>G)^ cell line by CRISPR/Cas9 combined with asymmetric ssODN. (A) Experimental diagram of generation of the *HBB* −28 (A>G) mutation cell line in K562; (B) The region around *HBB* −28 are targeted with two asymmetric sgRNAs and ssODN are provided along with CRISPR/Cas9 DNA cleavage to generate *HBB* −28 (A>G) mutation. *sgRNA1* and *sgRNA2* are complementary to the sense and antisense strands respectively. Mutation site is indicated with red color in the middle of sequence. PAM: protospacer adjacent motif (orange); (C) Identification of the −28 (A>G) mutation by sanger sequencing. Expected mutation is shown in the red rectangle; (D) Expression of *HBB* mRNA determined by qPCR in K562^−28 (A>G)^ and K562; (E) Expression of *HBB* mRNA determined by ELASA with benzidine staining in K562^−28^ and K562.

### Transcriptome analysis of K562^−28 (A>G)^ and K562 before erythrogenic differentiation

Many K562^−28 (A>G)^ cells appear to have irregular morphology and spontaneous cell death compared to the wt counterpart, suggesting loss of HBB expression by −28(A>G) mutation may compromise cell viability in normal culture condition (supplementary Fig.1). To understand how the mutation affects molecular functions of cells at transcriptional level, we conducted RNA-seq to analyze the transcriptome differences between the isogenic cell line of K562^−28 (A>G)^ and its control K562. Pairwise Pearson correlation analysis revealed high similarity between replicates from K562^−28 (A>G)^ and K562 cells, indicating high reproducibility of our data (Fig. 2A). Overall, the gene expression levels between K562^−28 (A>G)^ and wt cell are similar transcriptome-wide, suggesting that the mutation may affect (Fig. 2B). We conducted analysis of differentially expressed genes (DEG) and found 120 and 524 genes were consistently upregulated and downregulated in K562^−28 (A>G)^ compared to K562 (Figure 2C). To further explore the affected underlying biological functions, we conducted GO term, KEGG and Reactome analysis using those DEGs and found pathways of (cellular) response to hypoxia and response to (decreased) oxygen level were upregulated in K562^−28 (A>G)^ mutant cell line (Figure 2D). In consistent, hypoxia related genes such as *HMOX1, BMP7, GATA6, ESAM, RYR2* were upregulated in K562^−28 (A>G)^ (Figure 2C). Interestingly, PI3K-Akt signaling pathway that is important for erythrocyte differentiation ^[18]^, was downregulated in K562^−28 (A>G)^ (Fig. 2D). To further explore the core regulators, we performed interaction assay to predict transcription factors (TFs) that target upregulated genes in K562^−28 (A>G)^. In agreement with previous results, we observed *GATA* family, *HOXD10* and *SPIC*, which are well-known regulators of erythroid differentiation and hypoxia [ref], were the core regulators for upregulated genes in K562^−28 (A>G)^ (Fig. 2E). The regulators and their corresponsive target genes were listed and genes related to hypoxia response were labelled in red (Fig. 2F). Taken together, these data suggested hypoxia response was upregulated in K562^−28 (A>G)^ and the core regulators were GATA family, HOXD10 and SPIC.

**Figure 2.**
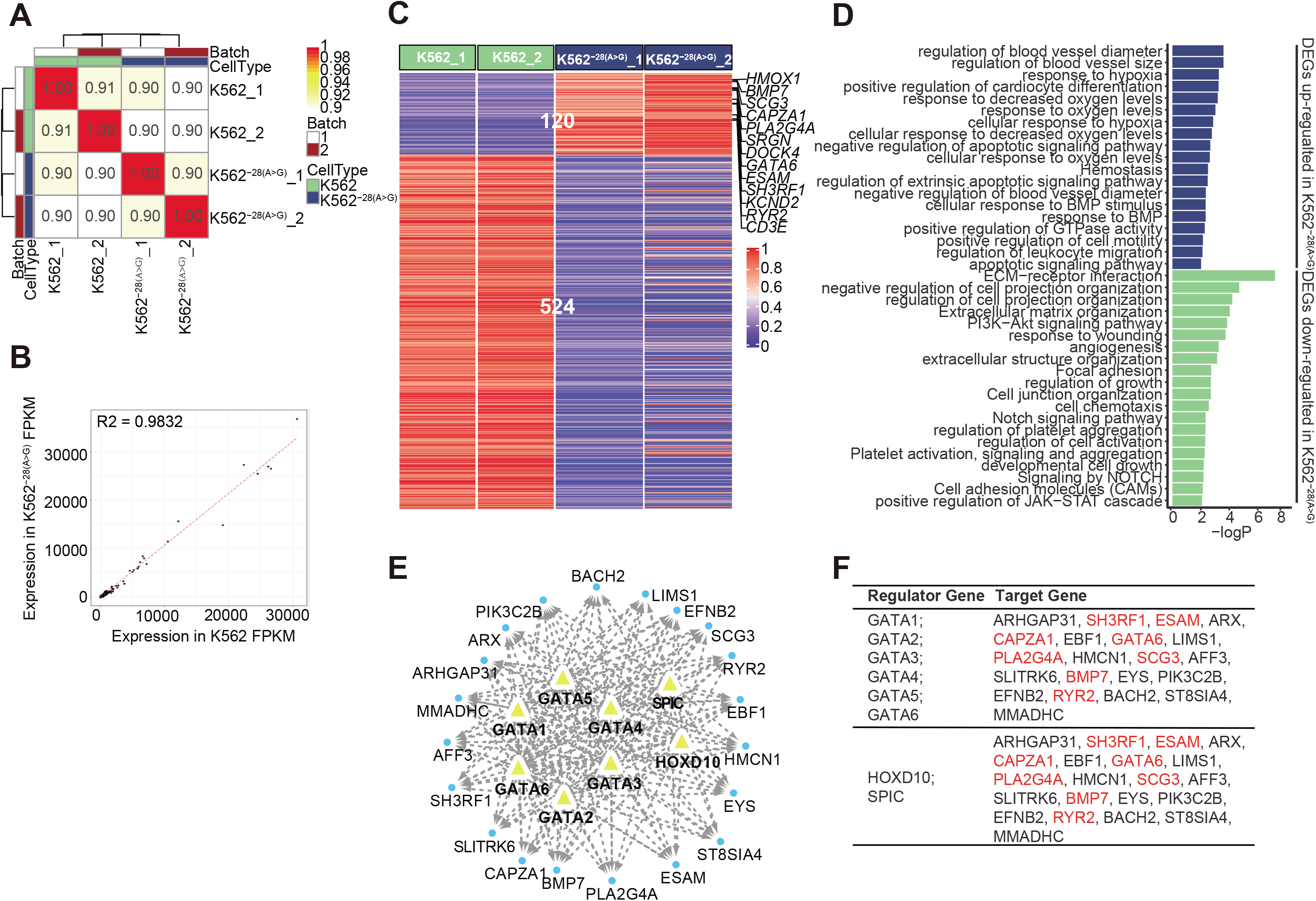
Transcriptome analysis of K562 and K562^−28 (A>G)^ cell lines before erythrogenic differentiation. (A) Pairwise Pearson correlations are represented in matrix between two K562 and two K562^−28 (A>G)^ samples before differentiation; (B) The correlation between representative K562^−28 (A>G)^ and K562 sample before differentiation; (C) Heatmap shows the differentially expressed genes (DEGs) between K562 and K562^−28 (A>G)^ cell lines; genes related to hypoxia were showed on right (D) Enrichment analysis of GO, KEGG and Reactome pathways based on DEGs in K562^−28 (A>G)^ before differentiation; (E) The predicted interaction map between transcription factors (TF) in DEGs of K562^−28 (A>G)^ before differentiation; (F) The regulator genes and their corresponsive target genes. The hypoxia gene were showed in red.

### Transcriptome analysis of co-regulated genes in K562^−28 (A>G)^ and K562 cell lines after erythroid differentiation

In order to investigate the effect of *HBB* −28 (A>G) mutation during erythroid differentiation, we induced erythroid differentiation and then performed genome-wide RNA-Seq in K562 and K562^−28 (A>G)^ mutant cell line. Pairwise Pearson correlations represented in matrix indicated that the isogenic cell lines showed high similarities up to 90% and the samples were clustered together depending on their differentiation conditions (Fig. 3A). The expression of 1385 genes were consistently upregulated in differentiated K562^−28 (A>G)^ and K562 when compared to undifferentiated samples (Fig. 3B) and these DEGs were used to conduct the GO term, KEGG and Reactome analysis. Multiple signaling pathways were activated in both K562^−28 (A>G)^ and K562 (Fig. 3C), including PI3K-Akt, MAPK and ERK pathways that have been reported to be activated during erythroid differentiation ^[18–21]^. In addition, signal pathways such as cell adhesion, pluripotency of stem cells, platelet activation and Notch pathway, were also co-activated in differentiated samples (Fig. 3C), suggesting our induction process was successful in both cell lines. *HBB* belongs to globin gene family mainly including *HBA* (α-globin), *HBG* (γ-globin) and *HBE* (ε-globin). *HBB, HBG* and *HBE* are β-globin-like and all capable of forming tetramer with a-globin. They are located within a gene cluster on chromosome 11 and their expression are coordinated by the same locus control region (LCR) with other regulatory DNA elements ^[22]^. In β–thalassemia patients, *HBG* may be upregulated to compensate the loss of *HBB*. To study the compensatory gene expression of globin genes in K562^−28 (A>G)^ after differentiation, the expression of *HBB, HBA* and *HBG* was analyzed by RNA-sequencing in isogenic cell lines of K562^−28 (A>G)^ and K562. As expected, Integrative Genomics Viewer (IGV) analysis showed that *HBB* expression was induced in K562 but undetectable in K562^−28 (A>G)^ after differentiation (Fig. 3D). In contrast, expression of other globin genes was induced and detected in K562^−28 (A>G)^ after differentiation (Fig. 3E). we noticed that fold induction rate of *HBA1, HBA2, HBE1 and HBZ* were similar or slightly increased in K562^−28 (A>G)^-Dif when compared to those in K562-Dif. However, *HBG* expression in K562^−28 (A>G)^-Dif was generally lower than K562-Dif (Fig. 3F). The expression of *HBB* and *HBG* before and after differentiation were further confirmed by RT-PCR. In consistent, expression of *HBB* was undetectable in undifferentiated K562^−28 (A>G)^ and its expression level in K562^−28 (A>G)^-Dif after differentiation was negligible when compared to K562-Dif control (Fig. 3G). In agreement with RNA-seq results, expression level of *HBG* was increased in both cell lines after differentiation, but induction fold of *HBG* was lower in K562^−28 (A>G)^ than that in K562 (Fig. 3G and 3H). We further analyzed the expression of key transcription factors that are important for regulation of globin genes during erythrogenic differentiation. KLF1, a positive regulator of *HBG* expression, is downregulated in K562^−28 (A>G)^. Expression of other positive regulators, including GATA1, BGL3 and NFE2 ^[19, 23–25]^, were dramatically upregulated in K562 after differentiation, while those increasement were largely attenuated in K562^−28 (A>G)^. In contrast, negative regulator BCL11A was upregulated in K562^−28 (A>G)^ before differentiation (Fig. 3I). Those data suggested the erythrogenic differentiation was overall normal in K562^−28 (A>G)^, but *HBB* −28 (A>G) mutation may affect the *HBG* expression through a network of transcription factors.

**Figure 3.**
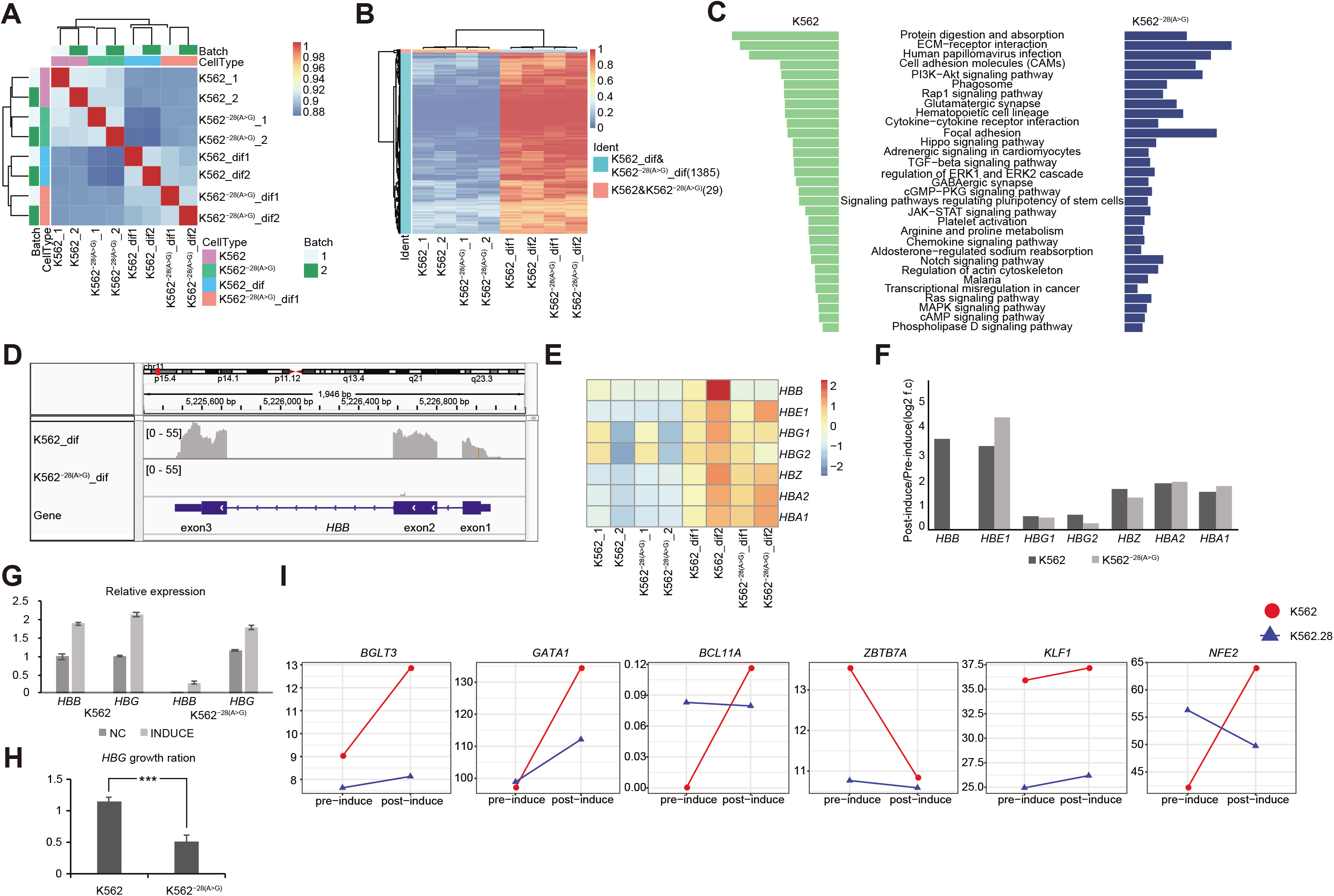
Transcriptome analysis of co-regulated genes, including HBB genes, in K562 and K562^−28 (A>G)^ cell lines after erythrogenic differentiation. (A) Pairwise Pearson correlations are represented in matrix between isogenic cell lines. The differentiated group are clustered together; (B) Heatmap shows the co-regulated genes in K562 and K562^−28 (A>G)^ after erythrogenic differentiation when compared to those before differentiation. (C) KEGG signaling pathways enriched in differentiated K562 and K562^−28 (A>G)^ when compared to their corresponding cell lines before differentiation; (D) The Integrative Genomics Viewer (IGV) shows the *HBB* gene expression in K562 and K562^−28 (A>G)^ cell lines; (E and F) Expression and change trends of globin genes determined by RNA-seq in the K562 and K562^−28 (A>G)^ cell lines before and after induction; (G and H) Expression and change trends of globin *HBB* and *HBG* determined by qPCR in the K562 and K562^−28 (A>G)^ cell lines before and after induction; (I) Expression of key transcription factors related to erythroid differentiation.

### Transcriptome analysis of differentially expressed genes (DEGs) in K562 and K562^−28 (A>G)^ cell lines after erythroid differentiation

To understand the effect of HBB −28 (A>G) mutation during erythrogenic differentiation, we identified the DEGs within K562^−28 (A>G)^-Dif and K562-Dif. The number of upregulated and downregulated DGEs in K562^−28 (A>G)^ were 158 and 740 respectively (Fig. 4A). With GO term, KEGG and Reactome analysis, we found up-regulated DEGs in K562^−28 (A>G)^-Dif were enriched in pathways related to stress-response and hematopoiesis disorder, such as regulation of apoptosis process, negative regulation of leukocyte activation, myeloid leukocyte cytokine production, negative regulation of blood coagulation, negative regulation of hemostasis, and negative regulation of hemopoiesis. Meanwhile, down-regulated DEGs in K562^−28 (A>G)^-Dif were enriched in oxygen related pathways, including oxygen transport, erythrocytes take up carbon dioxide and release oxygen and O2/CO2 exchange in erythrocytes (Fig. 4B). Reactome analysis of critical DEGs and their related KEGG pathways in differentiated K562^−28 (A>G)^-Dif was shown, and *PDE4D, TFPI* and *CA1, AQP1* which were related to hypoxia were found to be critical targets (Figure 4C and Supplementary table 1). Moreover, TF prediction revealed that JAZF1, MSI2, KDM4, HOX and ZNF family were core regulators for upregulated genes in K562^−28 (A>G)^-Dif (Fig. 4D), while SPIC and GATA family are core regulators for downregulated genes in K562^−28 (A>G)^-Dif (Fig. 4E). Interestingly, GATA3 was found to be the core regulator for both upregulated and downregulated genes (Fig. 4D and E). Taken together, dysregulated genes and pathways were present in K562^−28 (A>G)^ after differentiation and transcription factors, such as GATA, HOX and ZNF families, may play important roles as core regulators.

**Figure 4.**
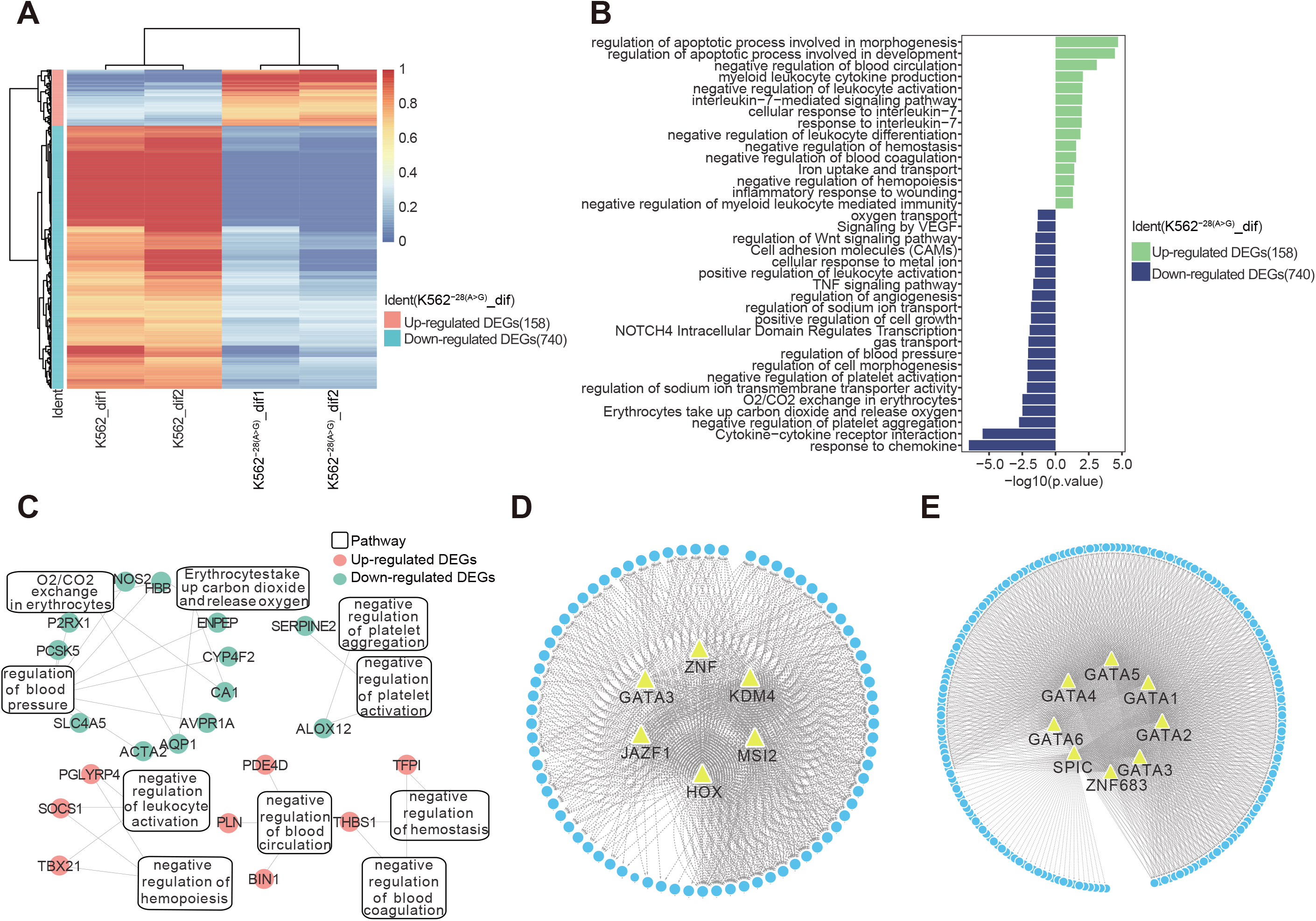
Transcriptome analysis of differentially expressed genes in K562 and K562^−28 (A>G)^ cell lines after erythrogenic differentiation. (A) Heatmap shows DEGs in K562^−28 (A>G)^ when compared to K562 after differentiation; (B) Signaling pathways analysis based on DEGs of K562^−28 (A>G)^ compared to K562 after differentiation; (C) Reactome analysis of critical DEGs and their related KEGG pathways in differentiated K562^−28 (A>G)^ when compared to its corresponsive K562 control. (D) the prediction of TFs in K562 after differentiation. (E) the prediction of TFs in K562^−28 (A>G)^ after differentiation.

### Reversion of observed abnormalities in corrected K562^−28cor^ cell line

As it took weeks for the generation of K562 mutant cells by Crispr/Cas9 and clone selection, it is possible that the diffrerential gene expression is due to secondary effects of the mutation and HBB loss, similar to studies of patient samples (ref). In order to determine the specificity of changed gene expression that attributes to the mutation, we corrected the *HBB* −28 (G>A) mutation in K562^−28 (A>G)^ by the same strategy of CRISPR/Cas9 combined with assODN (Fig. 5A and supplementary Fig.2), and named the corrected cell line as K562^−28(A>G)cor^. We analyzed the correlation of RNA-seq data between K562^−28(A^>^G)cor^, K562 and K562^−28 (A>G)^ and found expression profile of K562^−28(A>G)cor^ was closer to K562 rather than its precursor K562^−28 (A>G)^ (Fig. 5B). DEGs of isogenic cell lines were showed in the form of heat map and results indicated that upregulated and downregulated genes in K562^−28 (A>G)^ were reversed in K562^−28(A>G)cor^ (Fig. 5C). Through analysis of pathway enrichment, PI3K pathway and cell response to oxygen pathway were recovered in K562^−28(A>G)cor^ and the related genes, including *BMP7, EGR1* and *KCND2*, were showed in heatmap. The erythroid transcriptional factors such as *GATA1, KLF1* and *BCL11A*, were also recovered in K562^−28(A>G)cor^ (Fig. 5D and E and supplementary Fig. 5A and B). In summary, a large portion of abnormalities observed in K562^−28 (A>G)^ are reversed in corrected K562^−28(A^>^G)cor^, suggesting these phenotypes are specifically caused by mutation of *HBB* −28 (A>G).

**Figure 5.**
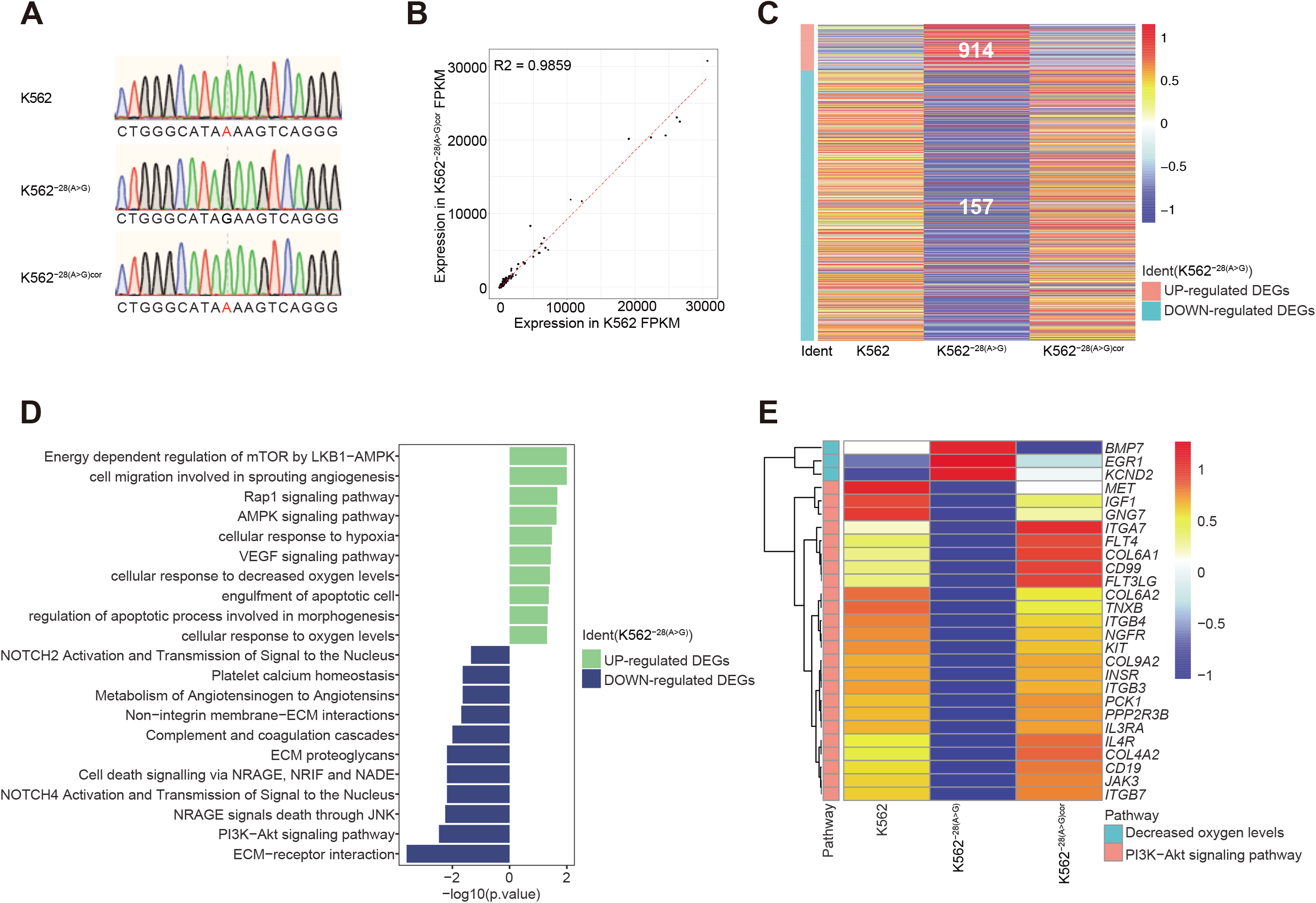
Genes and key pathways are reversed in mutation corrected K562^−28(A>G)cor^ cell line. (A) the identify of HBB gene in K562^−28(A>G)cor^ by sanger sequencing. (B) The correlations between the K562 and K562^−28(A>G)cor^; (C) Heatmap shows the up-regulated and down-regulated DEGs in K562, K562^−28 (A>G)^ and K562^−28(A>G)cor^; (D) Differentially regulated signaling pathways in K562^−28cor^ when compared to K562^−28 (A>G)^; (E) the recovery of genes relative to Key pathway and hypoxia.

## Discussion

In our study, we successfully generated the cell line K562^−28 (A>G)^ with *HBB* 28 (A>G) mutation using CRISPR/Cas9 and a 127bp assODN. This disease model of cell line was used to study how *HBB* −28 (A>G) mutation affected the cellular function on transcriptome level pre- and post-erythroid differentiation.

Our results showed *HBB* −28 (A>G) mutation prevented the transcription of *HBB* gene. Analysis of enriched pathways suggested PI3K pathway, as well as JAK-STAT pathway, which play important roles in the erythroid differentiation, were disrupted in K562^−28 (A>G)^ before erythroid differentiation. The PI3K-Akt signaling pathway is a significant pathway that control many cellular processes known as cell division, autophagy, survival, and differentiation [ref]. Moreover, the mutation activated the hypoxia pathway in undifferentiated K562^−28 (A>G)^. Many clinical manifestations observed in β–thalassemia is attributed to the chronic hypoxic environment due to pathologic erythrocyte production, and our data suggest hematopoietic precursors may also subject to oxidative stress before differentiation.

To induce erythroid differentiation, we chose the glutamine-minus medium with sodium butyrate, as hemoglobin synthesis was markedly induced using this condition with a differentiation efficiency of 11%~19% in K562 ^[26, 27]^. Consistent with previous reports, the differentiation efficiency of K562 in our study was nearly 12% (data not shown), indicating that our erythroid differentiation is effective. In agreement, the MAPK and ERK pathway was activated in both K562 and K562^−28 (A>G)^ (Fig. 2), a finding consistent with observations in previous studies ^[20, 28]^. Interestingly, PI3K-Akt signaling pathway was activated in K562^−28 (A>G)^ after induction, suggesting the defective PI3K pathway may be caused by lack of activators in undifferentiated K562^−28 (A>G)^. Other pathways, such as cell adhesion, pluripotency of stem cells, platelet activation and Notch pathway, were also co-activated in differentiated K562 and K562^−28 (A>G)^ samples, indicating mutation of *HBB* −28 (G>A) didn’t block the pathways required for differentiation. Nevertheless, in consistent with data from undifferentiated K562^−28 (A>G)^, oxygen related pathways were downregulated in differentiated K562^−28 (A>G)^ (Fig. 4). In both undifferentiated and differentiated conditions, SPIC and GATA families are predicted as core regulators (Fig.2 and Fig.4). The GATA family of transcription factors (GATA1-6) are essential for normal hematopoiesis and a multitude of other developmental processes ^[19, 29]^. *GATA-1* regulates terminal differentiation and function of erythroid which activates or represses erythroid-specific gene, such as β-globin locus-binding protein, and it might regulate the switch of fetal to adult haemoglobin in human ^[30]^.

Interestingly, increased expression of *GATA1* was largely attenuated in K562^−28 (A>G)^ during erythroid differentiation (Fig. 3), which may play a role for dysregulated oxygen related pathways.

As improving the levels of *HBG* in adults could partially reverse the severity of symptoms in sickle disease and β–thalassemia, it is important to understand the coordinated regulation between *HBB* and *HBG* ^[31–34]^. In this study, we noticed that fold induction of *HBG* was decreased in K562^−28 (A>G)^. *ZBTB7A* and *BCL11A* were two major repressors of *HBG* by directly bound the *HBG* gene promoters ^[24, 35–37]^. Expression of *ZBTB7A* was decreased during differentiation in K562, consistent with increased expression of *HBG*. However, the overall expression level was higher in K562 than that in K562^−28 (A>G)^, paradoxical with the results of decreased fold induction of *HBG* in mutant cell line. Although expression of *BCL11A* was lower in K562 when compared to 562^−28 (A>G)^ before differentiation, its expression increased dramatically during differentiation (Figure3I and supplementary Figure5). Collectively, our data suggest the expression of *HBG* was not only regulated by *ZBTB7A* and *BCL11A*, but may also be regulated by GATA proteins that also regulate *HBB*. However, the interaction requires further investigations.

Lastly, we not only used CRISPR/Cas9 to generate mutation but also corrected the mutation with the same strategy (Fig. 5A). On one aspect, reversed abnormalities in corrected cell line confirmed the specificities of these phenotypes. On another aspect, the efficient editing results indicate our gene editing strategy using assODN is powerful and provide means for gene editing treatment of *HBB* mutation with −28(A>G).

In summary, we show isogenic K562^−28 (A>G)^ cell line generated with CRISPR/Cas and assODN is a valuable model to evaluate the β-thalassemia with homozygous mutation of *HBB* −28 (A>G). It provides us with the first transcriptome data for mechanistic study on the effects of *HBB* −28 (A>G), and our disease model may also be used to assay the new therapies against β-thalassemia in the future.

## Data Availability

The data of our study have been deposited in the CNSA (https://db.cngb.org/cnsa/) of CNGBdb with accession code CNP0000981.

## ACKNOWLEDGEMENTS

We would like to thank Yanmei Deng, Jiaying Zhang and Wenwen Yao for helping single cell identification. We thank the Genome Synthesis and Editing Platform of the China National GeneBank for providing support on gene synthesis. We thank BGI colleagues for helping to output the high-quality data. This research was supported by National Natural Science Foundation of China (No.31970816 and 81903159), Guangdong Provincial Key Laboratory of Genome Read and Write (No. 2017B030301011) and Shenzhen Municipal Government of China (No.20170731162715261)

## AUTHOR CONTRIBUTIONS

Y.G., C.L. and X.Z conceived and designed the study. J.L performed experiment from L.G, T.L, G.D and Ouyang; Z.Z, and H.S performed bioinformatics analysis; J.L., Z.Z., Y.G and C.L. wrote the manuscript with the inputs from all authors.

## Methods

### Generation of β-thalassemia with-28 (A/G) cell line and corrected cell line sgRNA design and construction

two 20-bp sgRNAs were chosen containing the −28 (A/G) site, and the cutting site about 3 bp and 9bp before the mutation site respectively, the sequence we showed in Figure1B. The guide RNA oligonucleotides were synthesis by BGI and inserted into the gRNA cloning vector pSpCas9(BB)-2A-GFP (PX458) (Add gene 48138) according to the protocol provided by Zhang F’s protocol^[38]^.

Design of ss ODN repair templates: The 127-nt asymmetric ss ODN repair templates were designed by overlapped the CRISPR/Cas9 cleavage site with 36 bp on the PAM-distal side, and with a 91-bp extension on the PAM-proximal side of the break^[16, 17]^ (Figure1A and 1B) and were synthesized by BGI. Once the mutation of −28 (A/G) were mutated, the site was blocked and Cas9 could not cut this site again without anything changed. So, it is easy manipulate and seamless on the genome.

Gene editing: For gene targeting, 1*10^6 K562 cells were collected and resuspended in 100ul Buffer R at a final density of 1.0 × 107 cells/mL, electroporated with 3 mg each of the ss ODN, 1.5mg gRNA and Cas9 (Addgene48138) by Neon Transfection System(Invitrogen) Transfected cells were plated onto the prepared 6 well plates. After transfection 48h, we use flow cytometry to isolate the single cell in 96 well plates. about 1 week later, we Identify the single cell clones by PCR and sanger sequence. The resistant colonies were picked and expanded until we generated the expected cell line. primers are listed in Supplemental.

### Cell Cultures and differentiation

K562 cells obtained from American Tissue Culture Collection. the cells of K562 and K562^−28(A>G)^ were cultured in glutamine-minus RPMI 1640 medium(Gibco) with10% FBS in the presence of 1 mM sodium butyrate for 7 days to induce the erythroid differentiation, and then collected^[39]^. And the control cells were cultured in RPMI 1640 medium with 10% FBS and P/S antibiotics for 7 week at the same time.

### Real-time Reverse transcriptase (RT)-PCR analysis

Total RNA was isolated from the cells by the TRIzol™ Reagent (invitrogen). Single stranded cDNA was synthesized with the oligo(dT) primer using PrimeScript™ RT reagent Kit with gDNA Eraser (Takara), the obtained cDNAs were analyzed by real-time PCR, using indicated primers. The *HBB* primers were 5’-GCTCGGTGCCTTTAGTGATG −3’ (forward) and 5’-GCACACAGACCAGCACGTT −3’ (reverse); for *HBG*, 5’-GGAAGATGCTGGAGGAGAAACC −3’ (forward) and 5’-GTCAGCACCTTCTTGCCATGTG −3’ (reverse). The cycling conditions were: 95°C denaturation for 10 min, 95°C for 15 s, annealing and extension at 60°C for 40s, 40 cycles on the ABI step one machine.

### Comparison the hemoglobin expression

To comparison the hemoglobin expression of the uninduced cells (K562, K562^−28(A>G)^, K562^−28(A>G)cor^ dif) and induced cells. K562dif K562^−28(A>G)^ dif we used Tetramethylbenzidine(TMB) Elisa Kit (Invitrogen) .10^5^ cells from each of the test and control groups were resuspended in phosphate buffered saline (PBS) and mixed with equal volume of the Tetramethylbenzidine solution and then kept at room temperature for several mins. And then add double volume of Tetramethylbenzidine solution for stop solution. Brown-blue wells were regarded as positive and read at 450nm by Biotek epoch.Each treatment was performed in triplicate and each experiment was repeated for three times. Sample information in table1

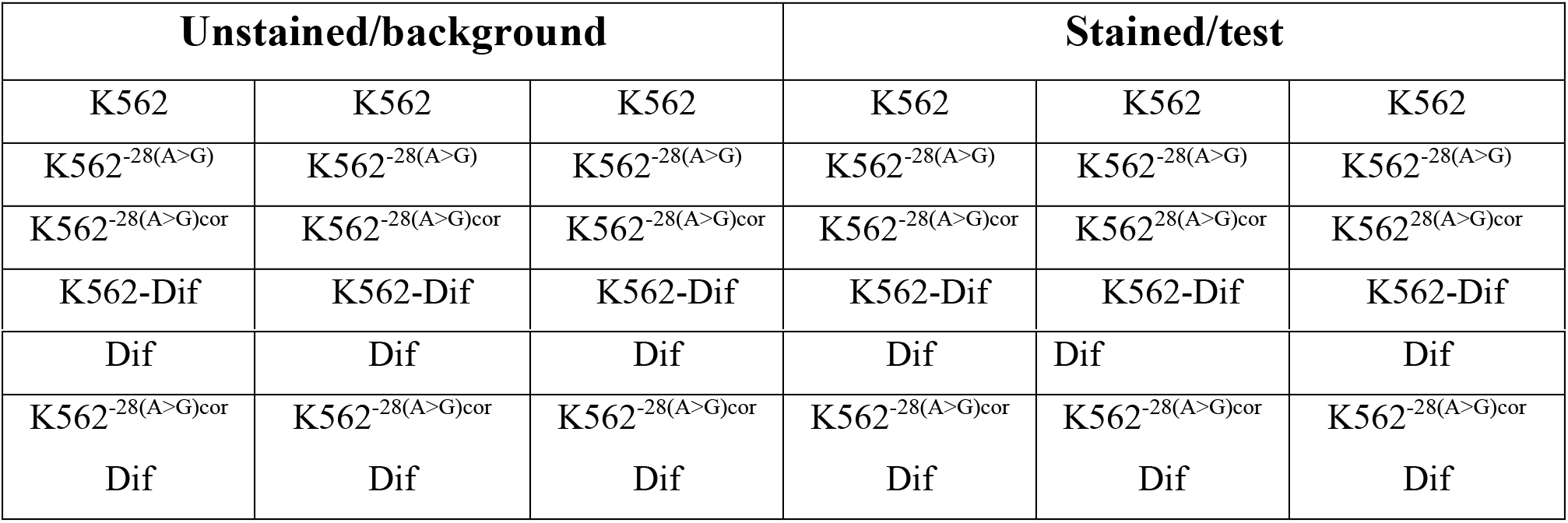

### Flow cytometry

We used CD235ab (Bio legend, USA) as the erythroid-specific surface marker which was determined by direct immunofluorescent staining with FITC-conjugated mouse monoclonal antibody. About 5 * 10^5^ cells were harvested and counted, resuspended in 100ul PBS with Human BD Fc Block™(BD) for about 10mins at room temperature, then add100 ul FITC-conjugated with cd235ab diluted by 1:10 and incubated at 4 °C for 30 min. Cells were washed with ice-cold PBS three times and The fluorescence was measured assayed by flow cytometry of BD Aria III. Unstained group was used as antibody stain control for each samples (K562, K562-28, K562 induced, K562-28 induced)

## RNA-Seq

### RNA-Seq analysis

There are exceeding 2 million 100 bp pair-ended reads in each sample. Prior to assembly, Raw reads were filtered by SOAPnuke ^[40]^ with the parameters “-l 15 -q 0.2 -n 0.05”. Reads were mapped to the human reference genome using HISAT2^[41]^, which included both the genome sequences (GRCh38.p12) and known reference sequence (RefSeq) transcripts. Finally, the 80% of reads were aligned to human genome.

We used Samtools (Trapnell et al., 2012)^[42]^ to sort and index the alignment files and Integrative Genomics Viewer (IGV) tool (Casero et al., 2015)^[43]^ was used to visualize the reads. The sorted binary sequence analysis file (BAM files) was also used to generate UCSC Browser tracks with a genome Coverage Bed from Bed Tools (Dennis et al., 2003)^[44]^. To this end, coverage files were normalized using the total signal for each sample.

The StringTie^[45]^ suite of tools was used to calculate and compare gene expression levels and normalized as FPKMs (Fragments Per Kilobase of transcript per Million mapped reads). Differential expression analysis was assessed at each sample by comparing the pre- and post-induction cells using EdgeR Bioconductor package^[46]^. Differentially expressed Genes (DEGs) were identified when the log_2_|fold change| >1 and the adjusted p-value <0.01.

### Gene ontology (GO) analysis

Gene ontology enrichment analysis of the gene sets was then performed on each sample using the cluster Profiler R package^[47]^, including KEGG^[48]^ pathways and Gene Ontology^[49]^ (http://www.geneontology.org/), which were collected in Molecular Signatures Database (MSigDB)^[50]^, Meanwhile, DAVID (https://david.ncifcrf.gov/) and Metascape were utilized to assess the enrichment of functional categories (GO and KEGG) of the DEGs^[51]^.

### Transcription factor prediction

To compare the expression levels of transcription regulator genes between pre-induced and post-induced K562 cells, we collected a comprehensive transcription factor annotation from AnimalTFDB 3.0^[52]^ and iRegulon^[53]^, and the results visualized using Cytoscape^[54]^.

## List of abbreviations

DMEM: Dulbecco’s modified Eagle’s medium
HBB: **Hemoglobin, Beta**
FCS: fetal calf serum
BSA: bovine serum albumin
assODN: asymmetric single strand oligo DNA nucleotides
K562^−28(A>G)^: HBB-28(A>G) cell line
K562^−28(A>G)cor^: Corrected HBB-28(G>A) cell line
Dif: Erythroid differentiation

