## supplemental figures for "Transcriptome analyses of β-thalassemia -28 (A>G) mutation using isogenic cell models generated by CRISPR/Cas9 and asymmetric single-stranded oligodeoxynucleotides (assODN)"

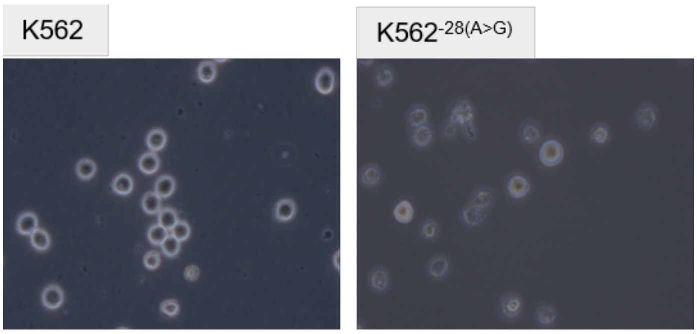


Figure1: the cell shape and morphology of K562 and K562 $^{(-28(A>G)}$

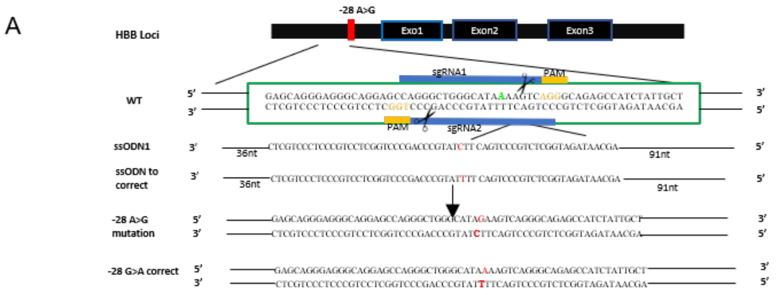


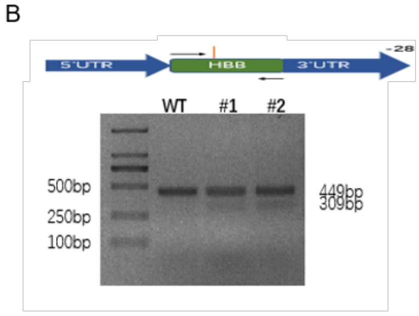


C

| | Sequencing number | Indel number | Mutation correct |
|---|---|---|---|
| Mutation | 86 | 8 (9.3%) | 1 (1.16%) |
| Correction | 60 | 55 (91%) | 3 (5%) |

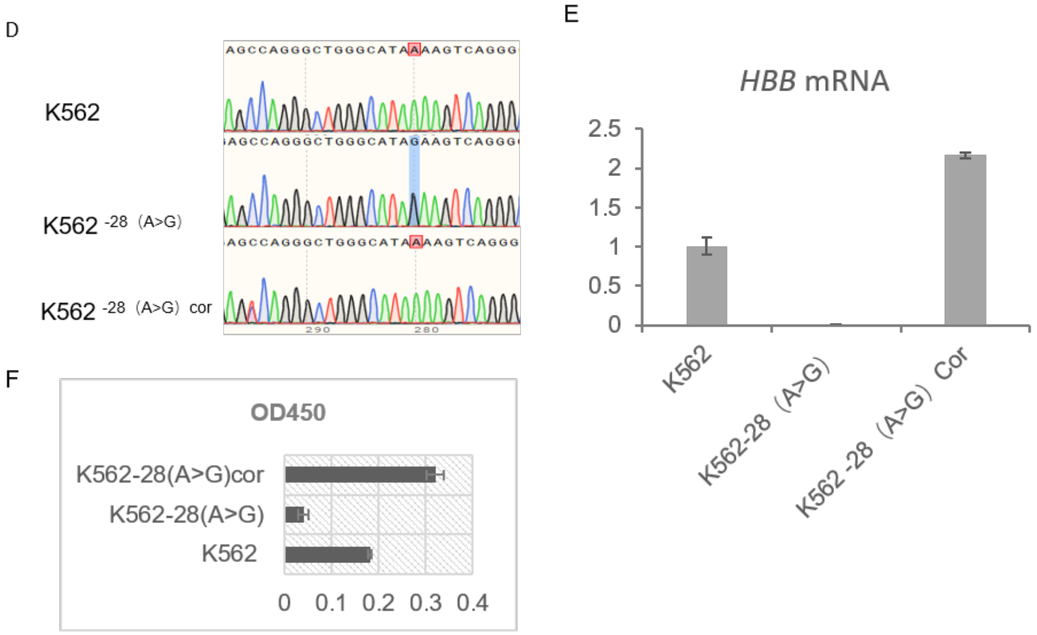


Figure2: Generation of K562-28 cell line by asymmetry ssODN combined CRISPR/Cas9

A:The region around HBB -28 are targeted with two asymmetric sgRNAs and ssODN are provided along with CRISPR/Cas9 DNA cleavage to generate HBB -28 (A>G) mutation. sgRNA1 and sgRNA2 are complementary to the sense and antisense strands respectively. Mutation site is indicated with red color in the middle of sequence. PAM: protospacer adjacent motif (orange)

B: gRNA activity assay #1,#2 represent gRNA1 and gRNA2.

C: the efficiency of the editing of isogenic cell lines.

D: Sanger sequencing identify the mutation and correction SNP of -28 site. Expected mutations showed in the red rectangle.

E: qPCR identify the expression of *HBB* in the three cell lines.

F: Determination the hemoglobin of isogenic lines by TMB

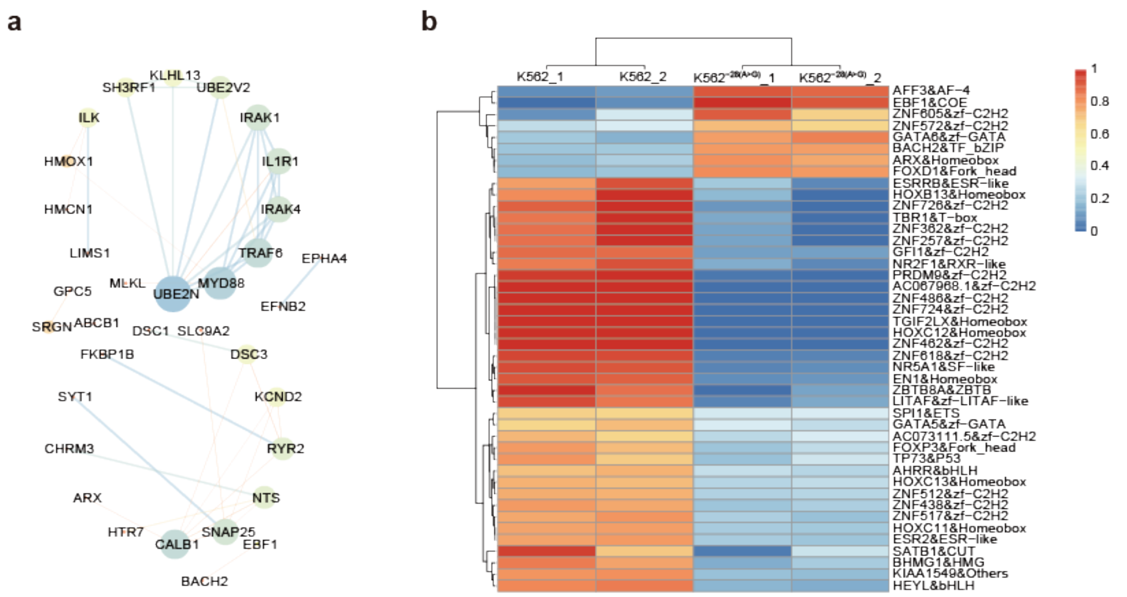


**Figure S3**

Figure3: the PPI network of hypoxia gene and TF prediction

A: The PPI network of hypoxia genes such *as HMOX1 ,SRGN.*

B: Heatmap reporting scaled expression of predicted transcriptional regulators in K562-28 and K562 cell lines before induction.

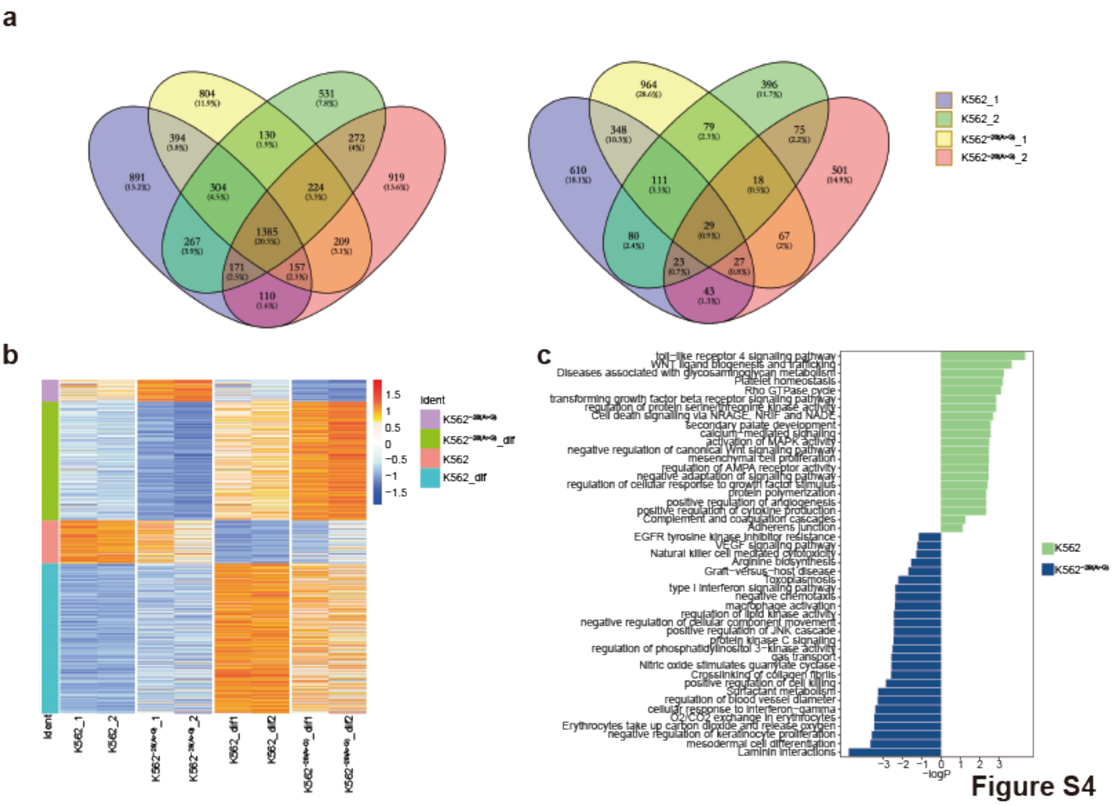


**Figure S4**

Figure4: the different DEGs and pathways of K562 and K562-28 pre and post induction

A: The DGEs of K562 and K562-28 induction process. The left and right are up-regulated and down-regulated DEGs after induction in K562 and K562-28, respectively.

B: Heatmap reporting scaled, imputed expression for the different DEGs in pre- and post-induction K562 and K562-28 cell lines .

C: Differentially expressed signaling pathways after induction in isogenic cell lines.

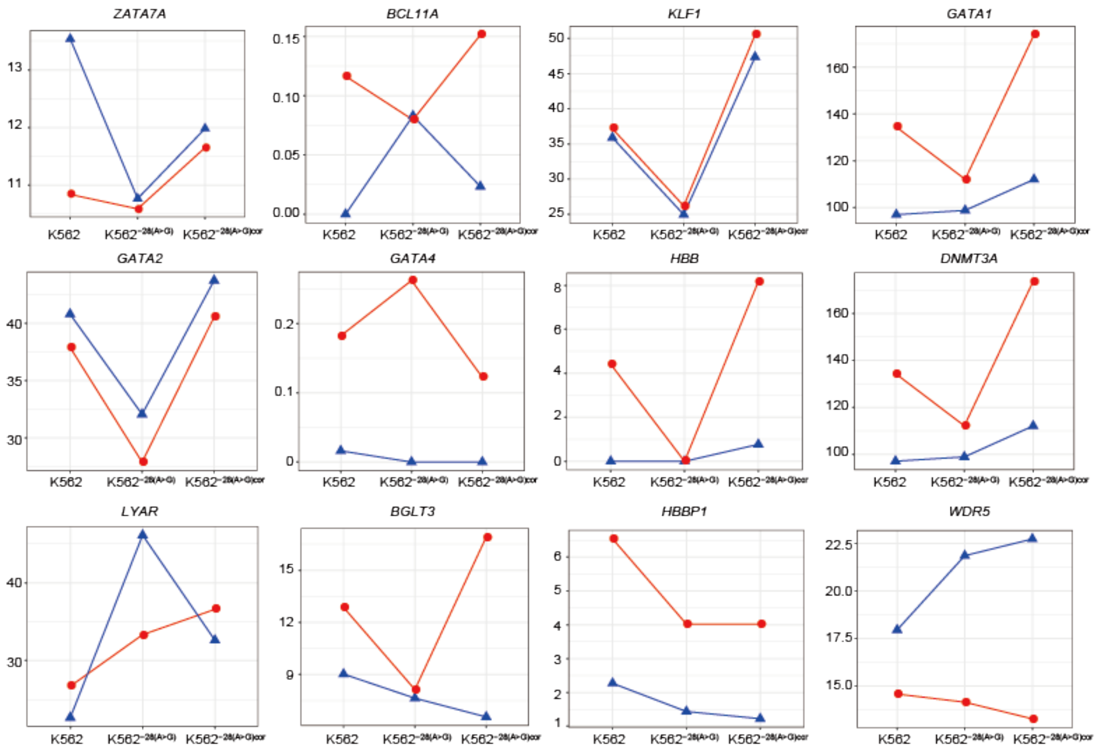


**Figure S5**

Figure5: the transcriptional factors expression in isogenic cell lines pre and post differentiation

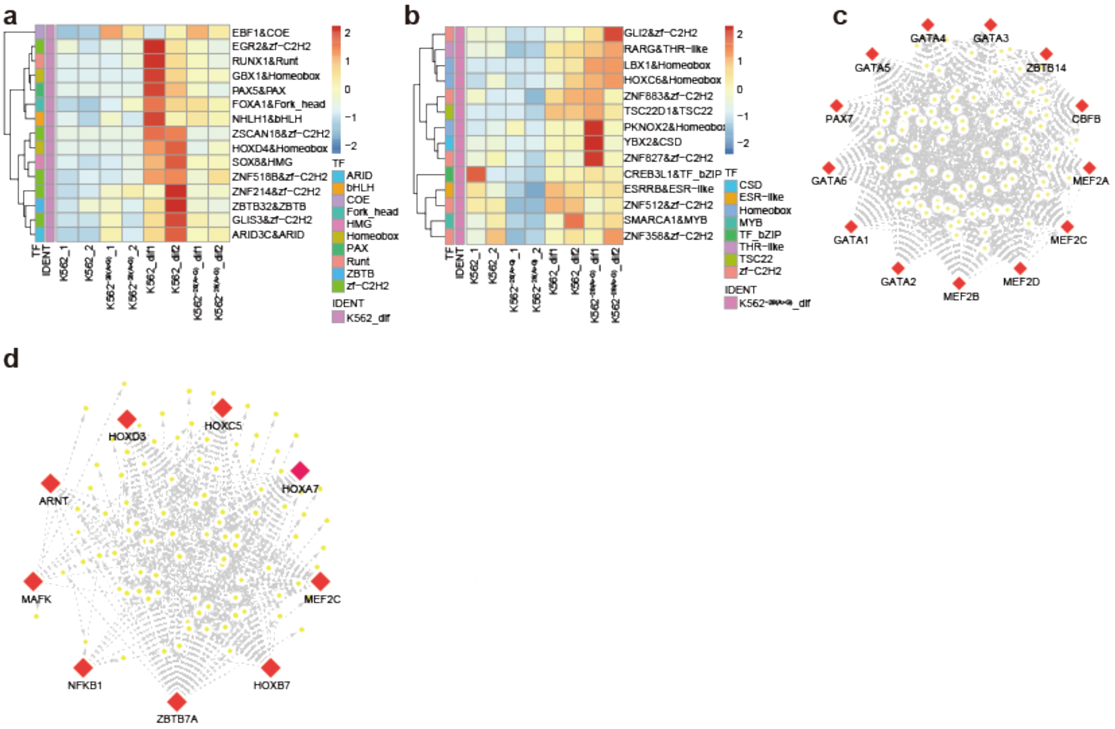


**Figure S6**

**Figure 6: Heatmap and TF predictition of K562 and $K562^{-28(A>G)}$**

A: Heatmap showing the high expression TFs in K562.Dif Rows represent up-regulated genes. Columns represent individual samples.

B: Heatmap showing the high expression TFs in $K562^{-28(A>G)}$ Dif. Rows represent up-regulated genes. Columns represent individual sample.

C: The predicted motifs and TFs highly expressed in Sample K562 Dif.

D: The predicted motifs and TFs highly expressed in Sample **$K562^{-28(A>G)}$** Dif.

| **Pathways** | **Gene** | **-Log$_{10}$(Pvalue)** |
|---|---|---|
| negative regulation of blood circulation (upregulated) | PDE4D, BIN1, PLN | 3.11 |
| negative regulation of leukocyte activation (upregulated) | TBX21, HMOX1, PGLYRP4, SOCS1 | 2.02 |
| negative regulation of blood coagulation (upregulated) | TFPI, THBS1 | 1.55 |
| negative regulation of hemostasis (upregulated) | TFPI, THBS1 | 1.55 |
| negative regulation of hemopoiesis (upregulated) | TBX21, PGLYRP4, SOCS1 | 1.40 |
| negative regulation of platelet aggregation (downregulated) | ALOX12, SERPINE2, UBASH3B | 2.74 |
| Erythrocytes take up carbon dioxide and release oxygen (downregulated) | CA1, AQP1, HBB | 2.49 |
| O2/CO2 exchange in erythrocytes (downregulated) | CA1, AQP1, HBB | 2.49 |
| negative regulation of platelet activation (downregulated) | ALOX12, SERPINE2, UBASH3B | 2.10 |
| regulation of blood pressure (downregulated) | NOS2, PCSK5, ACTA2, P2RX1, ENPEP, AVPR1A, CYP4F2, SLC4A5, SUCNR1, HBB | 2.07 |

Table 1: the key pathway and its genes in **K562**$^{-28(A>G)}$ Dif

| TAIR_ID | SYMBOL | Expression in K562(FPKM) | Expression in **K562**$^{-28(A>G)}$ | Expression in **K562**$^{-28(A>G)oor}$ |
|---|---|---|---|---|
| ENSG00000198712.1 | MT-CO2 | 30692.64844 | 36744.03516 | 30770.25977 |
| ENSG00000198727.2 | MT-CYB | 6165.318359 | 7140.681641 | 5964.253418 |
| ENSG00000198763.3 | MT-ND2 | 10497.41895 | 11490.30371 | 11902.92871 |
| ENSG00000198804.2 | MT-CO1 | 24538.55078 | 25426.05859 | 20618.97852 |
| ENSG00000198840.2 | MT-ND3 | 5910.532227 | 6203.662598 | 4681.167969 |
| ENSG00000198886.2 | MT-ND4 | 26198.11133 | 26931.85742 | 23103.125 |
| ENSG00000198888.2 | MT-ND1 | 12149.9541 | 15624.04492 | 11654.20117 |
| ENSG00000198899.2 | MT-ATP6 | 6574.290527 | 8397.078125 | 6698.809082 |
| ENSG00000198938.2 | MT-CO3 | 22358.1543 | 27259.70703 | 20360.51367 |
| ENSG00000210082.2 | MT-RNR2 | 19094.99414 | 14788.95996 | 20154.99023 |
| ENSG00000210140.1 | MT-TC | 7179.018066 | 6783.394531 | 5114.338867 |
| ENSG00000210144.1 | MT-TY | 5791.18457 | 5859.809082 | 4686.176758 |
| ENSG00000212907.2 | MT-ND4L | 26618.31641 | 26450.43555 | 22554.38477 |
| ENSG00000225630.1 | MTND2P28 | 5174.062012 | 5560.36377 | 4965.740234 |
| ENSG00000248527.1 | MTATP6P1 | 6778.824707 | 8012.343262 | 5417.968262 |

**table 2**

Table 2: the highly expression genes associated with mitochondria in **K562**$^{-28(A>G)}$ Dif
